# Somatosensory Realignment Following Single and Dual Force Field Adaptation

**DOI:** 10.1101/2025.07.08.663774

**Authors:** Dylan Zangakis, Aaron L Wong, Amanda S. Therrien

## Abstract

Evidence that adaptive motor learning coincides with a realignment of somatosensory perception has led to hypotheses that a shared mechanism underlies both processes. This implies that these two phenomena should exhibit similar properties. However, studies of somatosensory realignment with visuomotor adaptation have shown mixed support, possibly due to a confounding coactivation of sensory prediction errors and multisensory integration. While the former is thought to drive adaptation, both processes may contribute to somatosensory realignment. Here, we examined somatosensory realignment following force field adaptation, which is not confounded by multisensory integration. Across two experiments, we tested whether somatosensory realignment mimics three canonical properties of adaptation in this paradigm. Our first experiment examined whether sensory realignment (for the perception of movement or static position) correlated with adaptation across individuals, and generalized beyond the trained reach direction. The results showed that force field adaptation coincided with a selective realignment of somatosensory perception of movement in the direction of the perturbing force, but this realignment did not correlate with the magnitude of adaptation or generalize beyond the reach direction of the adaptation task. In a second experiment, we tested whether context-dependent dual adaptation to opposing force field perturbations coincides with a context-dependent dual realignment of somatosensory perception. The results showed no evidence of context-dependent somatosensory realignment after dual adaptation. Overall, our results indicate that somatosensory realignment and adaptation exhibit different properties and are therefore unlikely to rely on the same underlying mechanism, although realignment does display some coherence with the nature of the perturbation.

**NEW AND NOTEWORTHY:** This study is the first to demonstrate a dissociation in the realignment of somatosensory perceptions of static position and movement following adaptation to novel forces in the upper limb. By assessing whether perceptual realignment following force field adaptation exhibits two canonical properties of this type of motor learning – i.e., generalization to nearby movement directions and context dependence – this study constitutes a key test of theories positing a shared mechanism underlying the motor and perceptual processes.

## INTRODUCTION

Motor adaptation describes the process by which an individual learns to update their motor commands to counter a consistent perturbation (Krakauer et al. 2019; Shadmehr et al. 2010). When initially applied, the perturbation induces movement errors that signal a discrepancy between the actual sensory consequences of a motor command and what was predicted – i.e., a sensory prediction error. Through a combination of cognitive and sensorimotor mechanisms, the direction and magnitude of errors are used to update predictions and, in turn, the motor commands based on them to account for the perturbation. This updating process continues until movement errors are minimized. Successful adaptation can be verified if unexpected removal of the perturbation results in movement errors opposite to those caused by the initial introduction of the perturbation (i.e., after-effects).

In addition to motor output, a growing body of work shows that adaptation also changes somatosensation – the perception of body position and movement in the absence of vision (Proske and Gandevia 2012). The majority of evidence for this comes from studies of adaptation to visuomotor rotation perturbations, where the visual consequences of a movement are rotated at a consistent angle from the movement (and somatosensation) of the fingertip during target-directed reaching (Cunningham 1989; Krakauer et al. 2000). Somatosensory perception is often assessed by asking individuals to report the location of the unseen fingertip. Following adaptation to a visuomotor rotation perturbation, the perception of fingertip location tends to be shifted in the direction of the visual offset (Barkley et al. 2014; Block and Bastian 2011, 2012; Clayton et al. 2014; Cressman and Henriques 2009, 2011; Harris 1963, 1965; Henriques and Cressman 2012; Maksimovic et al. 2020; Modchalingam et al. 2019; Nourouzpour et al. 2015; Ostry and Gribble 2016; Ruttle et al. 2022; Salomonczyk et al. 2011; Tsay et al. 2021).

The co-occurrence of somatosensory realignment with motor adaptation has led to hypotheses that these two phenomena are mediated by a common underlying mechanism (Rossi et al. 2021; Tsay et al. 2022). Regardless of its exact nature, a shared mechanism implies that somatosensory realignment should display similar properties to motor adaptation; in particular, the extent of adaptation and sensory realignment should be correlated. Yet, evidence supporting this idea has been mixed. Studies of visuomotor rotation adaptation during reaching have shown that somatosensory realignment lags behind motor changes (Zbib et al. 2016) and displays a different pattern of generalization (Cressman and Henriques 2015). Furthermore, there is substantial variance across studies in the degree of correlation between the extent of adaptation to visuomotor rotation perturbations and the magnitude of the concomitant somatosensory realignment.

It is difficult to compare motor adaptation and somatosensory realignment in visuomotor paradigms because the perturbation is thought to elicit two processes (Block and Bastian 2012). The imposition of novel visual consequences of movement commands triggers sensory prediction errors. Simultaneously, the misalignment of visual and somatosensory feedback of limb position triggers multisensory integration (Babu et al. 2023; Block and Liu 2023; Hsiao et al. 2022; Mostafa et al. 2019; Wali et al. 2023). While sensory prediction errors may drive both adaptation and sensory realignment, multisensory integration is primarily associated with the latter. Differences in the relative contribution of these processes to adaptation and somatosensory realignment may explain some of the discrepant properties noted in previous work. To avoid this issue, it is useful to compare adaptation and somatosensory realignment using a force field paradigm that can trigger sensory prediction errors without misaligning feedback from different sensory modalities. In these paradigms, participants must learn to counter predictable perturbing forces that push the target-directed reaches off of a straight-line trajectory (Shadmehr and Mussa-Ivaldi 1994).

Fewer studies have examined the somatosensory changes that occur with adaptation of reaching movements to novel force fields (Haith et al. 2008; Mattar et al. 2013; Ohashi et al. 2019; Ostry et al. 2010). Using various methods for assaying somatosensory perception, these studies show an overall biasing in the direction of the perturbing force that tends to correlate with the degree of adaptation. While these results are suggestive of a common underlying mechanism driving the two processes, somatosensory realignment still lags behind motor changes in force field adaptation (Mattar et al. 2013), and little is known about the degree to which somatosensory realignment mimics other properties of adaptation.

As noted above, force field perturbations affect the execution of reaches, rather than misaligning visual and somatosensory feedback of limb position. Adaptation to an analogous perturbation in the lower extremities has been shown to selectively realign the somatosensory perception of limb movement, but not limb position. That is, split-belt treadmill adaptation, which alters leg speed during walking, coincided with realignment of somatosensory perceptions of leg speed and the step length covered in a given time (Sombric et al. 2019; Vazquez et al. 2015). Yet, individuals showed no realignment of perceived foot position during standing or walking (Vazquez et al. 2015). A distinction between the senses of movement and position is well supported by prior studies of somatosensory perception (Brown et al. 2003a, 2003b; Proske 2019; Wong et al. 2024). However, no study to date has carefully dissociated whether the somatosensory realignment following adaptation in the upper extremity differently affects the perceptions of limb movement and limb position; failing to distinguish between these sensations may also contribute to the inconsistent evidence for a correlation between motor adaptation and sensory realignment. Given the nature of force field perturbations, it is reasonable to hypothesize that adaptation might coincide with a selective realignment of the perception of limb movement during reaching.

An additional, well-studied property of motor adaptation is the extent to which learning generalizes to other movement directions. Force field adaptation is known to exhibit a cosine-like generalization function, in which the newly adapted movement pattern is seen only for movement directions close to the one trained, typically within 45 degrees (Donchin et al. 2003; Gandolfo et al. 1996; Gonzalez Castro et al. 2011; Huang and Shadmehr 2007; Mattar and Ostry 2007; Rezazadeh and Berniker 2019; Sainburg et al. 1999; Thoroughman and Shadmehr 2000). In line with this, one prior study demonstrated that somatosensory realignment following force field adaptation did not generalize to a displacement perpendicular (i.e., rotated 90 degrees) to the trained one (Haith et al. 2008). However, it is not currently known how the somatosensory realignment observed with force field adaptation generalizes to reach directions closer to the trained ones.

Finally, force field adaptation has been shown to exhibit context dependence (Addou et al. 2011; Cothros et al. 2009; Howard et al. 2013, 2015; Ingram et al. 2013; Osu et al. 2004; Richter et al. 2004; Sheahan et al. 2016; Wada et al. 2003). That is, individuals can concurrently adapt to opposing perturbations when each is paired with an appropriate context cue, such as a distinct follow-through movement cued by the presentation of an additional reaching target (Howard et al. 2015). Upon removal of the perturbations, the provision of each context cue can elicit opposing after-effects. It is currently unknown whether dual adaptation to opposing perturbations may concomitantly bias somatosensory perception in opposing directions, with each triggered by presentation of the respective context cue utilized during the adaptation task.

Here, we assessed the degree of correlation with adaptation (specific to the sense of position or movement), generalization, and context-dependence of somatosensory realignment following adaptation to velocity-dependent force fields in the upper extremity. The first experiment showed that adaptation to a single force field was followed by a systematic and selective realignment of the perception of limb movement so that it was biased in the direction of the force perturbation, but the extent of adaptation and sensory realignment was not correlated across individuals. This realignment was specific to the reach direction trained in the adaptation task and did not generalize to other reach directions. A second experiment showed no evidence of context-dependent somatosensory realignment following dual adaptation to opposing force-field perturbations. We discuss our results in terms of how they elucidate prior findings of somatosensory changes following force field adaptation and how they fit into prior studies of the distinction between perception and action.

## MATERIALS AND METHODS

### Participants

A total of 52 right-handed adult neurotypical individuals were recruited for this study. Thirty individuals participated in Experiment 1 and were randomly assigned to one of two groups, each experiencing either a clockwise or counterclockwise curl force perturbation (clockwise group: n=15, average age 25.1 years, standard deviation 4.2 years, 2 identified as male, 13 identified as female; counterclockwise group: n=15, average age 25.5 years, standard deviation 4.5 years, 3 identified as male, 12 identified as female). Twenty-two individuals participated in Experiment 2 (average age 26.6 years, standard deviation 6.3 years, 6 identified as male, 16 identified as female). One individual participated in both Experiments 1 and 2 (aged 23 years, identified as female). The sample sizes for each experiment were chosen to be comparable with previous studies using similar tasks (Mattar et al. 2013; Ohashi et al. 2019; Ostry et al. 2010). All participants were naïve to the purposes of this study and provided written informed consent. Experimental methods were approved by the Thomas Jefferson University Institutional Review Board, and participants were compensated for their participation at a fixed hourly payment ($20/hour).

### Apparatus

Both experiments were performed using a Kinarm Exoskeleton Lab (Kinarm, Canada), which restricted the motion of the arms to the horizontal plane. Hand, forearms, and upper arms were supported in troughs appropriately sized to participants’ arms, and the linkage lengths were adjusted to match the limb segment lengths of each participant to allow smooth motion at the shoulder and elbow. Direct vision of the arm was obstructed by a horizontal mirror, through which participants were shown targets and cursors in the same plane as their movement via an LCD monitor (60 Hz refresh rate). All movements of the arm and forces applied by the robot were recorded at 1kHz.

### Experiment 1 Procedure

In this experiment, participants learned to adapt their reaching movements to counter a velocity-dependent curl force that pushed their arm in either a clockwise or counterclockwise direction (Shadmehr and Mussa-Ivaldi 1994). We assessed their perceptions of hand position (at reach endpoints) and the direction of hand movement during reaches before and after completion of the adaptation task (Ohashi et al. 2019). A schematic of the experimental paradigm is shown in Figure 1a.

**Figure 1.**
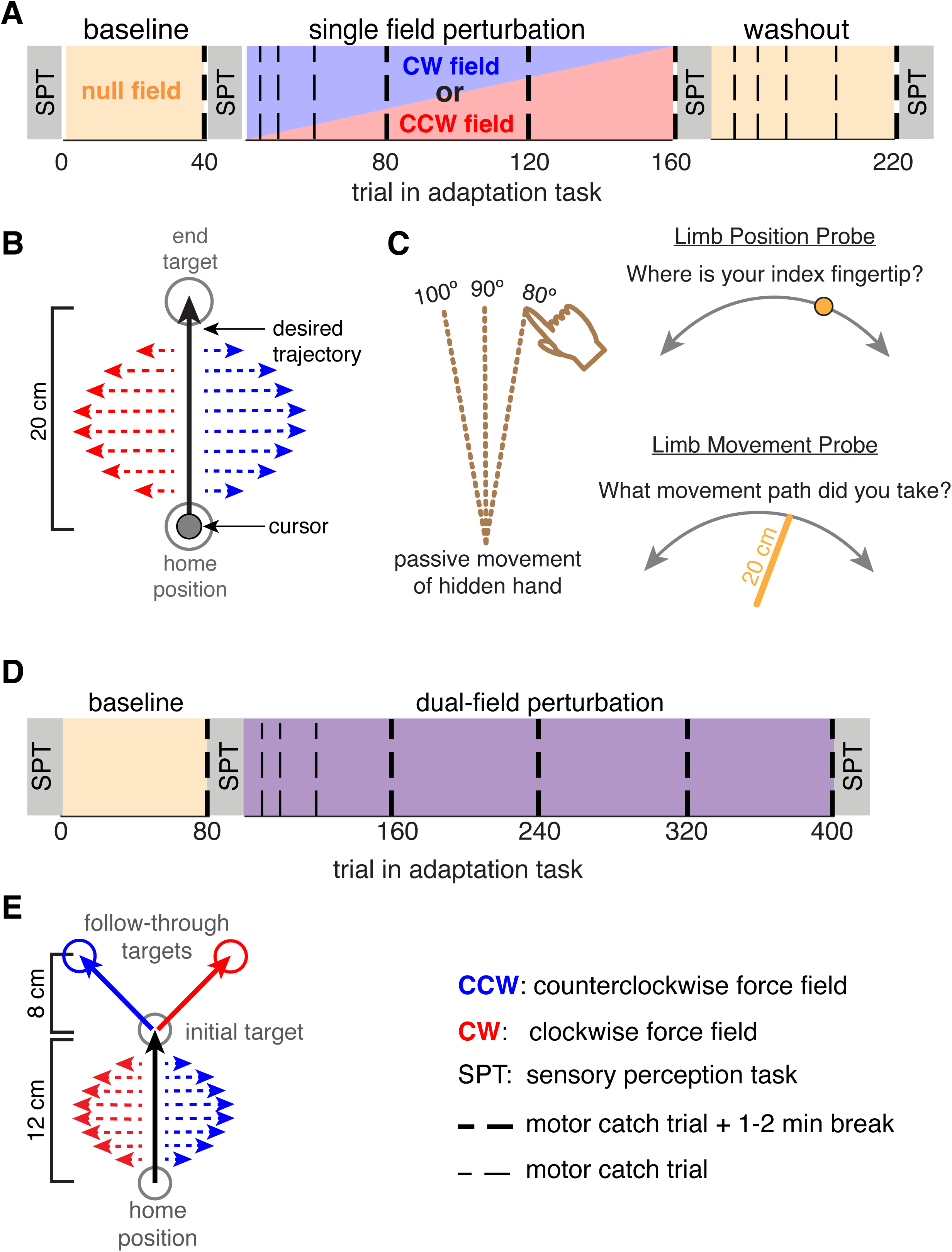
Schematic of the paradigms and experimental tasks in Experiments 1 and 2. **A.** The paradigm for Experiment 1. Participants completed a force field adaptation task that began with a baseline phase of reaching in a null field. Following the baseline phase, participants completed a perturbation phase of reaching in a force field. The perturbation phase was then followed by a washout phase where the force field was removed, and participants were again placed in a null field. Throughout the adaptation task, participants experience motor catch trials (vertical dashed lines), in which reaches were constrained to a force channel. Additionally, participants completed a series of sensory perception task blocks. Sensory perception task blocks were completed before and after the baseline phase, after the perturbation phase, and after the washout phase of the adaptation task. **B.** Schematic of the force field perturbation in Experiment 1. Participants completed 20 cm long reaches from a home position to a single end-target. A visual cursor indicated the position of the index fingertip throughout reaching movements. In the perturbation phase of the adaptation task, reaches were performed in a velocity-dependent curl field that pushed the hand in either a clockwise or counterclockwise direction. **C.** Schematic of the sensory perception task. Participants were passively moved through 20 cm reaching movements at one of 3 possible angles with the hand and arm hidden from view. In limb position probe trials, the hand was held at the reach endpoint, and participants were instructed to use a rotating dial to align the position of a visual cursor with the perceived current location of their index fingertip. In limb movement probe trials, participants were moved out and back along the reach path and instructed to use the rotating dial to adjust the angle of a line until it matched the perceived movement path. **D.** The paradigm for Experiment 2. Participants completed a dual-adaptation task that began with a baseline phase of reaching in a null field. The baseline phase was followed by a perturbation phase in which two force field perturbations were applied. The presentation of each force field varied randomly from trial to trial. Throughout the adaptation task, participants encountered motor catch trials (vertical dashed lines). In addition to the adaptation task, participants completed sensory perception task blocks at three points: before and after the baseline phase and after the perturbation phase of the adaptation task. **E.** Schematic of the dual force field perturbation. Participants made reaching movements from a home position that passed through a single initial target and followed through to one of two possible end targets. A velocity-dependent curl field was applied during the initial portion of the reach, the direction of which was mapped to the follow-through target.

#### Force Field Adaptation Task

Participants made 20 cm reaching movements with their dominant arm from a home position (1 cm radius circle located anterior to the sagittal midline) to an end target (1 cm radius circle located 20 cm directly anterior to the home position; Figure 1b). Visual feedback was provided throughout the task via a cursor aligned to the right index fingertip (0.5 cm radius circle). Each trial began with the participant holding their index fingertip in the home position. After a 500 ms delay, the end target appeared, cueing participants to begin their reach. Participants were instructed to reach in a straight line from the home position to the end target. Once the fingertip was held in the end target for 500 ms, the target changed color from blue to green, and an audible tone was played to signal the trial’s end. The end target was then extinguished, and participants could return their hand to the home position to begin the next trial. To encourage participants not to slow down while reaching, the words “Too Slow” appeared at the top of the experimental display if reaches took longer than 1200 ms to complete.

Participants completed a total of 232 reaching trials (Fig. 1a). The task was divided into three experimental phases. During the Baseline phase (reaching trials 1-41), participants performed their reaches in a null field (i.e., no forces were applied to the arm). During the Perturbation phase (reaching trials 42-167), a velocity-dependent force field perturbation was applied as follows:

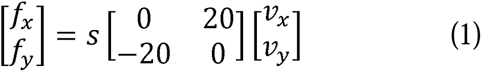

Where *x* and *y* are lateral and sagittal directions, *f_x_* and *f_y_* are the commanded forces to the hand in *N*, *v_x_* and *v_y_* are velocities of the hand in m/s, and *s* is the direction of the force field. A value of *s* = 1 corresponded to a clockwise force field, resulting in forces that deflected the hand to the right, while *s* = −1 corresponded to a counterclockwise force field that deflected the hand to the left (Figure 1b). Following the Perturbation phase, participants completed a Washout phase (reaching trials 168-232) in which the force field was removed, and reaches were once again made in a null field. Throughout the adaptation task, participants experienced 13 motor catch trials, in which the force field was unexpectedly removed and replaced with a 0.5 cm wide force channel that constrained reaches to a straight line. Motor catch trials occurred on the following reaching trials: 41, 47, 53, 64, 85, 126, 167, 173, 179, 190, 211, 232.

#### Sensory Perception Task

This task had participants report the sensed limb position or direction of movement following passive displacement of the right arm by the Kinarm robot (Figure 1c). The visual cursor representing the right fingertip position was removed for these trials. As in the adaptation task, the Kinarm’s display screen blocked the view of the arms and hands. On each trial, the robot passively moved the hand outward along a 20 cm reach path that began from the same home position used in the adaptation task and stopped at one of three possible invisible end positions, as shown in Figure 1c: (1) the position of the end target in the adaptation task (the 90-degree target), (2) a position 10-degrees clockwise to the end target (the 80-degree target), and (3) a position 10-degree counterclockwise to the end target (the 100-degree target). In Limb Position Probe trials, the hand was held at the end position, and a prompt appeared at the top of the screen asking, “Where is your fingertip?” A cursor (a 0.5 cm radius circle) also appeared on the screen at a random position along an invisible arc centered on the 90-degree target and spanning 160 degrees of the workspace. Participants were instructed to use their left hand to rotate a dial that was fixed to the Kinarm’s left hand tray to adjust the position of the cursor along the arc until they perceived it to match the position of their right index fingertip. Each notch of the dial rotated the cursor position by 0.5 degrees. In Limb Movement Probe trials, the robot returned the hand to the home position, reversing the outward trajectory. Participants were then presented with a prompt asking, “What movement path did you take?” At the same time, a line appeared on the screen that spanned the 20 cm radial distance between the home position and a random point along the invisible arc. Participants were instructed to turn the dial with their left hand to adjust the angle of the line until they perceived it to match the path traveled by their right index fingertip during the passive movement. In both trial types, participants verbally reported when they had finished adjusting the cursor/line. The experimenter then triggered the start of the next trial.

Sensory perception task blocks were interspersed throughout the force field adaptation task (Figure 1a), occurring before and after the baseline phase (pre- and post-baseline blocks, respectively), following the perturbation phase (post-perturbation block), and following the washout phase (post-washout block). Each block consisted of 36 trials (i.e., 6 repetitions of each passive reach direction for both the limb position and limb movement probes). Limb position and limb movement probes were completed in mini blocks of 3 trials, in which the three passive reach directions were randomized before switching the probe type.

### Experiment 2 Procedure

In this experiment, participants learned to concurrently adapt their reaching movements to counter two distinct and opposing velocity-dependent curl forces that pushed their arm in either a clockwise or counterclockwise direction. We assessed their perceptions of hand position (at reach endpoints) and the direction of hand movement during reaches before and after completion of the adaptation task to investigate the effects of dual adaptation on sensory recalibration. A schematic of the experimental paradigm is shown in Figure 1d.

#### Dual force field adaptation task

Participants had to simultaneously adapt to two opposing force fields by pairing each field to a distinct follow-through movement, as in prior work (Howard et al. 2015). Specifically, participants made 20 cm reaching movements with their dominant arm. Reaches began at a home position (1 cm radius circle located anterior to the sagittal midline) and passed through an initial target (1 cm radius circle located 12 cm directly anterior to the home position) before continuing on to slice through one of two possible follow-through targets (1 cm radius circles located at +/- 45-degree angles from the initial target at a radial distance of 8 cm; Figure 1e). As in Experiment 1, visual feedback of the right index fingertip position was provided throughout the task via an aligned cursor (0.5 cm radius circle). Each trial began with the participants holding their right index fingertip in the home position. After a 500 ms delay, two targets were presented on the display – the initial target and one of the two possible follow-through targets – along with text instructions to “Plan your reach.” During this phase, participants were instructed to remain in the home position and plan a reach that moved their index fingertip in a straight line from the home position to the initial target, then to continue on and slice through the second target. After a 700 ms delay, an auditory tone, along with a text display reading, “Go,” cued participants to enact the reach they had planned. Participants were required to accurately cross both the initial and follow-through targets in proper succession to complete the trial. This ensured that participants maintained accuracy in reaching target endpoints throughout all reaching trials performed. As in Experiment 1, the words “Too Slow” appeared at the top of the experimental display when reaches took longer than 1200 ms to complete.

Participants completed a total of 416 reaches. The task was divided into two experimental phases. During the baseline phase (trials 1-82), participants performed their reaches in a null field (i.e., no forces were applied to the arm). During the Perturbation phase (trials 83-416), a velocity-dependent force field that followed equation 1 was applied during the first portion of the reaching movement, when participants moved their index fingertip from the home position to the initial target. The force field was removed for the second portion of the movement when participants moved their index fingertip to slice through the indicated follow-through target. Here, the force field direction (and the value of *s* in equation 1) was mapped to the follow-through target such that a counterclockwise field was always paired with a follow-through movement to the right of the midline, and a clockwise field was always paired with a follow-through movement to the left of the midline (Figure 1e). Throughout the task, participants also performed a series of motor catch trials, which proceeded similarly to other trials, but the force field was unexpectedly removed and replaced with a 0.5 cm wide force channel that constrained the initial portion of reaches to a straight line. Motor catch trials occurred on the following trials: 81, 82, 93, 94, 105, 106, 127, 128, 169, 170, 251, 252, 333, 334, 415, 416.

#### Sensory Perception Task

Participants performed a modified version of the Sensory Perception Task from Experiment 1. Here, sensory perception trials began similarly to reaching trials. Participants began with their right index fingertip in the home position and would first be presented with two reach targets (the initial and a follow-through target) along with the text prompt to “Plan your reach.” However, participants would then see a text prompt reading “Sensory Test” instead of the go-signal, which would be followed by the robot passively moving the right arm through a 12 cm long reaching movement without visual cursor feedback, corresponding to the distance between the home position and initial reaching target in the dual force field adaptation task. Both limb position and limb movement probes then proceeded as they did in Experiment 1, with participants using a dial to report their perceived finger endpoint position or movement path. Based on the results of Experiment 1, which showed that the sensory effects of force field adaptation were localized to the trained movement direction, sensory perception trials only tested movements to the initial reaching target (i.e., at 90 degrees). To encourage participants to actually plan to counter the appropriate force field in each trial, a small number of adaptation trials were randomly intermixed during the sensory perception test, such that participants could not anticipate whether they were going to experience a sensory test trial or an adaptation trial.

### Data Analysis

All kinematic and kinetic analyses were performed using custom-written scripts in Matlab (MathWorks), and statistical analysis was performed using SPSS v30 (IBM).

In both of the force field adaptation tasks, the mean perpendicular deviation (MPD) for each reaching trial was computed by first constructing a line connecting the start and end points of each reach trajectory, then averaging the perpendicular deviation of each point along the trajectory to that line (Mattar et al. 2013; Ohashi et al. 2019; Ostry et al. 2010). In both tasks, the start of each reach was determined as the point at which the index fingertip left the home position. In Experiment 1, the end of the reach was determined as the position of the index fingertip after it had been in the end target for 500 ms. In Experiment 2, the end of the reach was determined as the position of the index fingertip when it entered the initial target. Like the targets, the y-axis was aligned to the sagittal midline, so negative MPDs corresponded to leftward (i.e., counterclockwise) deviations, while positive MPDs corresponded to rightward (i.e., clockwise) deviations. We quantified the peak velocity of each reach as an additional measure of movement kinematics. Finally, we measured the force compensation exhibited on motor catch trials as the correlation between the force exerted on the channel walls in the x-axis and the ideal force profile computed using the exhibited velocity (Howard et al. 2015). The correlation was multiplied by 100 and expressed as a percentage.

For each sensory perception task, we quantified reports as the signed difference between the angle of the actual end position or movement path induced by the robot and the angle reported by subjects using the cursor (for limb position probes) or line (for limb movement probes). Following this convention, a negative value corresponded to an error counterclockwise to the target, while a positive value corresponded to a clockwise error.

As both experiments aimed to quantify perceptual recalibration following motor adaptation, our first statistical analysis assessed which participants showed significant learning in each adaptation task. For each participant in Experiment 1, we used paired-samples t-tests to compare reach MPDs between the first 20 trials and the final 20 trials of the perturbation. Participants who showed a significant decrease (for the CW group) or increase (for the CCW group) in reach MPD were considered learners. In the dual adaptation task of Experiment 2, reach MPDs were more variable trial-to-trial, so paired-samples t-tests were used to compare the first and final 40 trials of the perturbation phase for each field separately. To be considered a learner in Experiment 2, the change in reach MPD had to be significant for both force fields. Participants who did not exhibit significant learning were not included in the final data analysis. This resulted in data from 5 of the 30 participants in Experiment 1 (2 from the CW group, 3 from the CCW group) and 7 of the 22 participants in Experiment 2 being discarded.

The primary statistical analysis of learners’ data was performed using linear mixed-effects regression models. All models were fit using restricted maximum likelihood methods and included subject-specific intercepts as random predictors. In all cases, the significance of fixed effects was tested using *F* tests on type III sums of squares with an alpha value of 0.05. The denominator degrees of freedom were estimated using Satterthwaite’s method. Post hoc analysis of significant interactions was conducted by estimating the marginal means for each level of factors in the interaction term, and then performing pairwise tests of simple effects with an alpha value of 0.05 adjusted for multiple comparisons using the Bonferroni method.

In experiment 1, both reach MPD and peak velocity were assessed as the dependent variables in separate models with fixed effects predictors of group (CW, CCW), phase (the first 20 trials of the perturbation, the final 20 trials of the perturbation, the first 20 trials of washout, and the final 20 trials of washout), and the group-by-phase interaction. As we needed to test whether both groups learned similarly, the reach MPD values of the CCW group were multiplied by −1 to correspond to the CW group to permit a valid comparison. Baseline perceptual reports were assessed as dependent variables in separate models for each target with fixed effects predictors of group (CW, CCW), phase (pre- and post-baseline reaching), probe type (limb position, limb movement), and all interaction terms. Post-adaptation and post-washout perceptual reports were normalized by subtracting the mean of the post-baseline reaching phase. The normalized reports were then assessed using separate models for each target with fixed effects predictors of group (CW, CCW), probe type (limb position, limb movement), and the group-by-probe interaction term. Normalized reports by the CCW group were not flipped, so as to highlight a difference in the direction of realignment between groups.

In experiment 2, reach MPD and peak velocity were assessed as the dependent variables in separate models with fixed effects predictors of field (CW, CCW), phase (the first 40 trials and the final 40 trials of the perturbation), and the field-by-phase interaction term. Baseline perceptual reports were assessed as dependent variables in a model with fixed effects predictors of field (CW, CCW), phase (pre- and post-baseline reaching), probe type (limb position, limb movement), and all interaction terms. As in experiment 1, post-perturbation perceptual reports were normalized by subtracting the mean of the post-baseline reaching phase. The normalized reports were then assessed as the dependent variable in a model with fixed effects predictors of field (CW, CCW), probe type (limb position, limb movement), and the field-by-probe interaction term. As was done for the Experiment 1 data, reach MPD values for the CCW force field were multiplied by −1, whereas the normalized perceptual reports were not flipped.

Our secondary statistical analysis involved additional tests of motor learning and changes in perceptual reports in both Experiments. As an additional test of adaptation, we analyzed the force compensation shown on motor catch trials. In Experiment 1, participants completed a single catch trial at 12 time points during the task. As the data here comprised a single observation per subject, we tested for a within-subject effect of time point using analysis of variance (ANOVA) with repeated measures (the alpha value was set to 0.05). Post hoc analyses were conducted using pairwise tests of simple effects with a Bonferroni correction for multiple comparisons. In Experiment 2, participants completed 2 catch trials, one for each context/field direction, at 8 time points during the adaptation task. As data here comprised two observations per subject, we tested for a fixed effect of time point using linear mixed-effects regression analysis following the same design and method for determining significance as noted above.

To further test whether the normalized perceptual reports post-perturbation (Experiments 1 and 2) and post-washout (Experiment 1) represented a significant change from baseline, we used one-sample t-tests to compare the distributions of normalized trial-level data to zero. In all cases where analysis of perceptual reports showed null results, we calculated the Bayes Factor (BF_01_) to quantify the strength of evidence for the null hypothesis over the alternative hypothesis. BF_01_ values are commonly interpreted as follows: 1 indicated no evidence for the null hypothesis, 1-3 indicated anecdotal evidence for the null hypothesis, 3-10 indicated moderate evidence for the null hypothesis, 10-30 indicated strong evidence for the null hypothesis, 30-100 indicated very strong evidence for the null hypothesis, and >100 indicated extremely strong evidence for the null hypothesis.

Finally, based on the results from the analyses described above, we performed some additional post hoc statistical tests. Pearson correlations were used to test the strength of the relationship between the change in reach MPD and the change in reach peak velocity over the perturbation phase for both Experiments 1 and 2. In Experiment 1, we also used linear mixed-effects regression and Pearson correlation to test the relationship between the change in reach MPD over the perturbation phase and the post-perturbation normalized limb movement perceptual reports.

## RESULTS

### Experiment 1

This experiment tested the specificity to movement and position perception and the generalization of somatosensory realignment following force field adaptation. Participants completed an adaptation task that had them reaching to a single target with their dominant arm using a robotic device. After an initial baseline phase, they had to learn to adapt their reaches to account for a perturbing force field that pushed their arm in either a clockwise or counterclockwise direction. Following exposure to this perturbation, they completed a washout phase where the force field was removed. Interspersed throughout the adaptation task, participants performed tests of sensory perception that assayed their sense of limb position following a passive movement, and their sense of the movement trajectory itself. Sensory perception was tested at three target angles: 90 degrees (corresponding to the movement performed during the force field adaptation task), and two generalization angles of 80 and 100 degrees.

#### Participants successfully adapted to the force field perturbation

Figure 2a shows the group mean time series of reach MPD values for both groups in the force field adaptation task. Statistical analysis of reach MPDs showed a main effect of phase (*F*(3,1969)=922.828, *p*<.001) that was driven by significantly greater MPD values in the early perturbation phase compared to all other phases of the task (all *p*<.001) – a result consistent with participants showing progressive adaptation to the force field perturbation. Reach MPD values in the early washout phase were also significantly different than the late washout phase (*p*<.001), indicating the presence of after-effects following the adaptation. There was no main effect of group (*F*(1,23.006)=1.670, *p*=.209), but the interaction between group and phase was significant (*F*(1,1969)=7.785, *p*<.001). Post hoc analysis showed the interaction was driven by differences in reach MPD between groups at two phases. In the late perturbation phase, the CW group showed greater reach MPDs, indicating that they learned slightly less than the CCW group (*p*=.033). Reach MPDs were also different between groups at the late washout phase, with the CCW group showing a more persistent after-effect than the CW group (*p*=.041).

**Figure 2.**
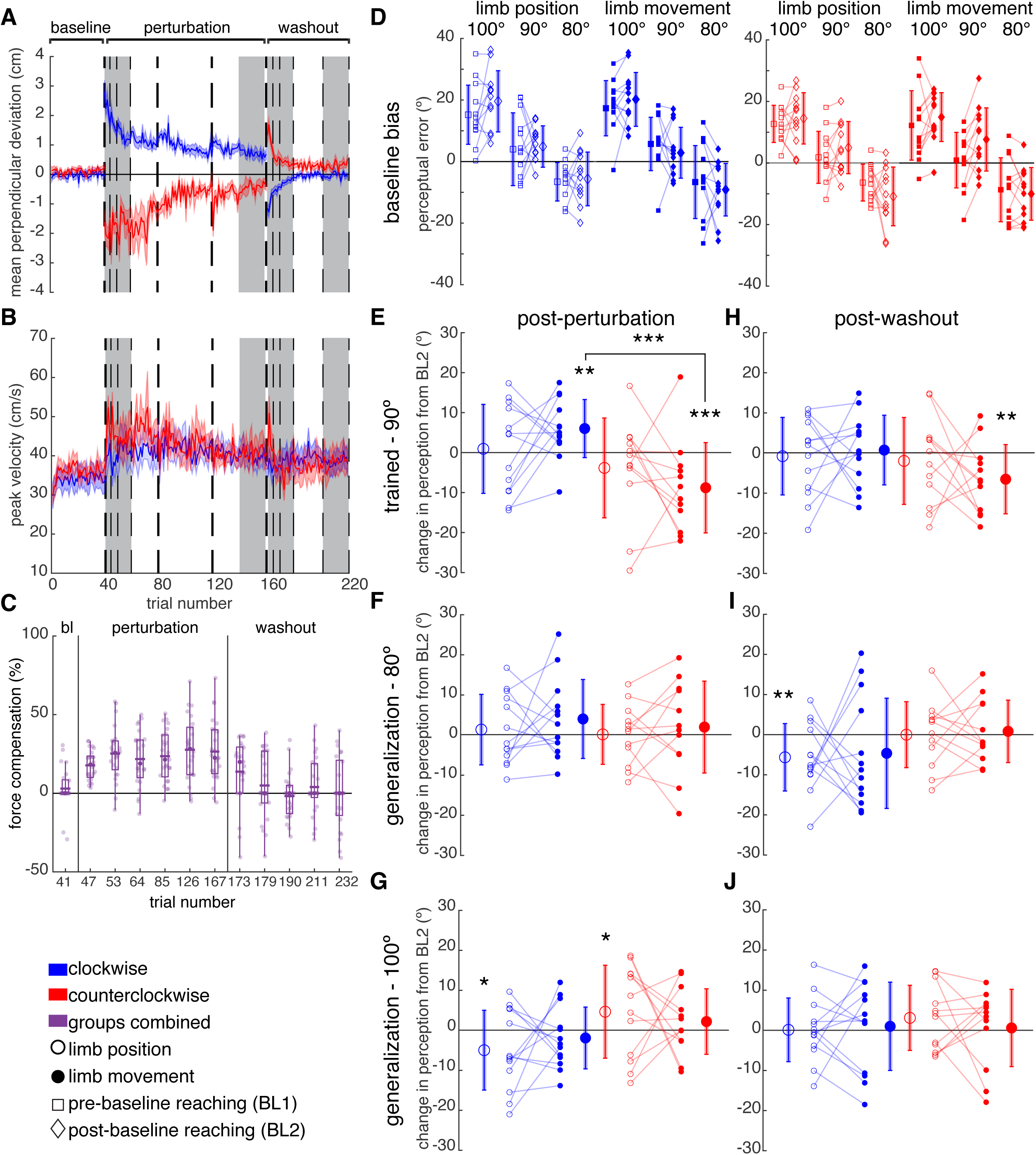
Experiment 1 results. **A-B.** Group mean time series of the mean perpendicular deviation values (**A**) and reach peak velocity (**B**) for the clockwise and counterclockwise perturbation groups. Vertical dashed lines depict motor catch trials. Thicker vertical lines denote points at which catch trials were followed by a 1-2 minute break. Grey shaded regions denote the early and late phases of the perturbation and washout periods. **C.** Force compensation on motor catch trials. Vertical boxes indicate the interquartile range, and whiskers indicate the data range excluding outliers. Horizontal lines show the distribution means, while solid markers indicate the median. Transparent markers show subject data. **D.** Perceptual errors exhibited as baseline by the CW (left panel) and CCW (right panel) groups. The report error is shown for each baseline test block within each target/movement angle for both perceptual probe types. **E-G.** The change in perceptual error between the sensory perception test blocks post-baseline reaching (BL2) and post-perturbation. Perceptual reports are shown for the trained target/trajectory angle at 90 degrees (**E**), as well as the generalization targets/trajectory angles at 80 (**F**) and 100 (**G**) degrees. **H-J.** The change in perceptual error between the sensory perception test blocks post-baseline reaching (BL2) and post-washout. Perceptual reports are again shown for the trained target/trajectory angle at 90 degrees (**H**), and the generalization targets/trajectory angles at 80 (**I**) and 100 (**J**) degrees. **D-J.** The smaller markers indicate individual subject means. Connecting lines depict the within-subject effects across testing phases (**D**) or perceptual probe types (**E**-**J**). The larger markers indicate group means, with error bars indicating standard deviation. *** *p*<.001, ** *p*<.01.

Since the force fields were velocity-dependent, any differences in reach velocity between groups or over task phases would indicate concomitant differences in the perturbation magnitude experienced by participants. Figure 2b shows the group mean time series of reach peak velocities over the adaptation task. Statistical analysis showed a significant effect of phase (*F*(1,1500.863)=33.849, *p*<.001), driven by a progressive slowing of reach velocity over the task. Early perturbation reach velocities were faster than all other task phases (all *p*<.001), and late perturbation reach velocities were also greater than both washout phases (both *p*<.001). Reach peak velocities were not significantly different between the early and late washout phases (*p*=1.000). There was no main effect of group (*F*(1,1991)=.212, *p*=.645), indicating that the CW and CCW groups behaved similarly overall. However, the group-by-phase interaction was significant (*F*(1,1500.863)=9.845, *p*<.001). Post hoc analysis of the interaction showed that the main effect of phase was largely driven by the CCW group, who showed all the same differences noted in the main effect (all *p*<.01). In contrast, the CW group showed no significant differences in peak velocity across phases (all *p*>.05). To determine whether the slowing of reach velocities could have confounded adaptation in the CCW group, we assessed the correlation between the changes in peak velocity and reach MPD. The result was not significant (CCW: *r*=.447, *p*=.145), suggesting that any slowing of reach velocities was unrelated to the degree of adaptation.

Analysis of the force compensation values showed a significant main effect of time point (*F* (11,264)=10.699, *p*<.001, Figure 2c). Post hoc analysis showed the effect was driven by force compensation at the end of baseline (i.e., trial 41) being significantly smaller than at each of the perturbation time points (all *p*<.05). The force compensation at the end of the perturbation phase (i.e., trial 167) was also significantly greater than that shown in the last 3 time points of the washout phase (i.e., trials 190, 211, and 232; all *p* <.05). Overall, these results provide additional evidence that participants adapted to the force field perturbation.

#### The perceptions of limb position and movement were not systematically different at baseline

Examining performance in the sensory perception task, it was important to first assess whether any differences existed between the groups at baseline and/or between the two baseline phases. Perceptual errors in the pre- and post-baseline reaching phases are shown in Figure 2d. Statistical analysis showed a significant interaction among group and phase at the 90-degree target (*F*(1,569)=4.107, *p*=.043), which was driven by a difference in the CCW group’s perceptual reports between the pre- and post-baseline phases (reports were significantly more positive overall in the second baseline phase compared to the first; *p*=.020). There was also a significant effect of phase at the 100-degree target (*F*(1,569)=4.604, *p*=.032), in which perceptual reports were significantly more positive in the second compared to the first baseline phase overall. We found no other significant effects, suggesting that perceptual reports did not systematically differ between probe types at baseline.

As a result of the differences observed between baseline phases, we normalized the post-perturbation and post-washout perceptual reports by subtracting the corresponding subject’s mean perceptual report from the second baseline (i.e., post-baseline) phase only. These normalized values were used to quantify the bias in perceptual reports following the force field perturbation and washout.

#### Force field adaptation selectively realigned the perception of limb movement

Figures 2e-g shows the normalized perceptual reports exhibited by the CW and CCW groups at each target tested in the post-perturbation phase. We first analyzed normalized reports at the 90-degree target (i.e., the one utilized during the adaptation task, Figure 2e). Statistical analysis showed a significant effect of group (*F*(1,23.002)=9.571, *p*=.005) and a group-by-probe interaction (*F*(1,273)=5.819, *p*=.017). The main effect of group was driven by the CW group showing an overall positive bias in their perceptual reports, whereas the CCW group showed an overall negative bias. However, post hoc analysis of the interaction showed that this group effect was asymmetric across probe types. The limb movement probe trials were significantly different between groups, with the CW group biased in the positive direction and the CCW group biased in the negative direction (*p*<.001). That is, both groups showed biases in limb movement perception that aligned with the direction of the force field perturbation. The groups did not significantly differ in limb position probe trials (*p*=.214, BF_01_=2.338). Within groups, the differences between probe types were not significant (CCW: *p*=.099, CW: *p*=.080). The overall effect of probe type was not significant (*F*(1,273)=.001, *p*=.980, BF_01_=29.731).

To determine whether the biases in limb movement perception at the 90-degree target comprised a significant shift from the baseline mean, we conducted one-sample t-tests to test whether the distributions of normalized perceptual reports differed from a zero-mean distribution. Indeed, the results were significant for both groups’ perceptions of limb movement (CCW: *t*(71)=-4.071, *p*<.001; CW: *t*(77)=2.823, *p*=.006), but not for the perceptions of limb position (CCW: *t*(71)=-1.751, *p*=.080, BF_01_=1.815; CW: *t*(77)=0.388, *p*=.699, BF_01_=7.455).

#### The realignment of limb movement perception did not generalize beyond the trained reach direction

The asymmetric effect found at the 90-degree target post-perturbation did not systematically generalize to either the 80- or 100-degree targets (Figures 2f and 2g, respectively). Statistical analysis showed no significant effects at the 80-degree target (group: *F*(1,23.004)=.304, *p*=.587, BF_01_=4.252; probe: *F*(1,273)=1.187, *p*=.277, BF_01_=4.380; group-by-phase: *F*(1,273)=.041, *p*=.840, BF_01_=19.497). Analysis of normalized perceptual reports at the 100-degree target showed a significant effect of group (*F*(1,23)=0.967, *p*=.004) in which the CW group showed an overall negative bias, while the CCW group showed a positive one. Yet, there was no effect of probe type (*F*(1,273)=.020, *p*=.888, BF_01_=10.060) and no interaction (*F*(1,273)=1.725, *p*=.190, BF_01_=7.731), indicating that the group behavior was consistent across perceptual probes.

One-sample t-tests also showed that none of the distributions at the 80-degree target differed significantly from a mean of zero (limb movement, CCW: *t*(71)=0.848, *p*=.399, BF_01_=5.466, CW: *t*(77)=1.828, *p*=.070, BF_01_=1.648; limb position, CCW: *t*(71)=0.039, *p*=.969, BF_01_=7.712, CW: *t*(77)=0.604, *p*=.524, BF_01_=6.581). At the 100-degree target, neither group showed a significant bias from zero in the perception of limb movement (CCW: *t*(71)=1.069, *p*=.289, BF_01_=4.469; CW: *t*(77)=-0.948, *p*=.346, BF_01_=5.204). However, the biases in limb position did reach significance (CCW: *t*(71)=2.097, *p*=.040; CW: *t*(77)=-2.342, *p*=.022).

#### The realignment of limb movement perception washed out with de-adaptation

Figures 2h-j show the normalized perceptual reports following the washout phase of the force field adaptation task. Analysis with the mixed-effects model showed no significant effects at the 90-degree target (group: *F*(1,23.048)=3.294, *p*=.083, BF_01_=1.262; probe: *F*(1,273)=.506, *p*=.478, BF_01_=6.448; group-by-probe: *F*(1,273)=2.024, *p*=.156, BF_01_=8.169; Fig. 2g). At the 80-degree target, there was a significant effect of group (*F*(1,23.028)=.209, *p*=.037; Fig. 2i) driven by the CW group showing a more negative bias than the CCW group overall. However, there was no effect of probe type (*F*(1,273)=.178, *p*=.673, BF_01_=36.998) or group-by-probe interaction (*F*(1,273)=.002, *p*=.968, BF_01_=7.621). At the 100-degree target, perceptual reports showed no significant effects (group: *F*(1,23.006)=.209, *p*=.652, BF_01_=6.607; probe: *F*(1,273)=.154, *p*=.695, BF_01_=7.423; group-by-probe: *F*(1,273)=.667, *p*=.415, BF_01_=52.650; Fig. 2j).

One-sample t-tests showed that the bias in limb movement perception exhibited by the CCW group post-washout remained significantly different from zero at the 90-degree (*t*(71)=-2.788, *p*=.007), but the CW group’s limb movement perception was not (*t*(77)=0.539, *p*=.591, BF_01_=6.969). Limb position perception at the 90-degree target remained comparable to a zero-mean distribution (CCW: *t*(71)=-0.715, *p*=.477, BF_01_=6.038; CW: *t*(77)=-0.177, *p*=.860, BF_01_=7.897). Analysis of the bias at the two generalization targets showed that the CW group’s limb position perception at the 80-degree target was significantly different from zero (*t*(77)=-2.721, *p*=.008), but there were no other significant effects (limb movement at 80-degrees, CCW: *t*(71)=0.455, *p*=.650, BF_01_=6.985, CW: *t*(77)=-1.905, *p*=.060, BF_01_=1.442; limb position at 80-degrees, CCW: *t*(71)=0.046, *p*=.963, BF_01_=7.710; limb movement at 100-degrees, CCW: *t*(77)=0.249, *p*=.804, BF_01_=7.491, CW: *t*(77)=0.469, *p*=.640, BF_01_=7.209; limb position at 100-degrees, CCW: *t*(71)=1.441, *p*=.154, BF_01_=2.876, CW: *t*(77)=0.060, *p*=.952, BF_01_=8.003).

#### Adaptation was not a significant predictor of biased limb movement perception post-perturbation

Analysis using a linear mixed-effects regression showed no significant relationship between the change in reach MPD over the perturbation phase and the post-perturbation normalized limb movement perception (*F*(24,50)=.482, *p*=.973, BF_01_=10.858). Pearson correlation analysis of the subject means for each measure was also not significant (*r*=-.035, *p*=.868). Thus, there was a systematic realignment of the perception of the trained movement following adaptation, but we did not find evidence that it was related to the degree of adaptation.

### Experiment 2

In a second experiment, we asked whether context-dependent adaptation also results in a context-dependent realignment of sensory perception. A new group of participants completed an adaptation task that had them making reaching movements from a home position to a single initial target and then following through to slice through one of two possible additional targets. After an initial baseline phase, a force field perturbation was applied, the direction of which switched randomly between CW and CCW from one trial to the next, but each direction was mapped to a particular follow-through target. In addition to the adaptation task, participants performed 3 blocks of sensory perception testing: before and after baseline reaching, and after the perturbation. Since Experiment 1 showed systematic realignment only at the 90-degree target, we probed the perceptions of limb movement and limb position only at this target.

#### Participants adapted to both force field directions

Figure 3a shows the group mean time series of reach MPD values exhibited during the context-dependent adaptation task. The results showed a significant effect of phase (*F*(1,2388)=622.034, *p*<.001), which was driven by greater MPD values in the first, compared to the final, 40 trials of the perturbation phase. There was no effect of field (*F*(1,2388)=.009, *p*=.924) or a phase-by-field interaction (*F*(1,2388)=2.58, *p*=.142), suggesting that participants showed comparable performance across the two force field directions.

**Figure 3.**
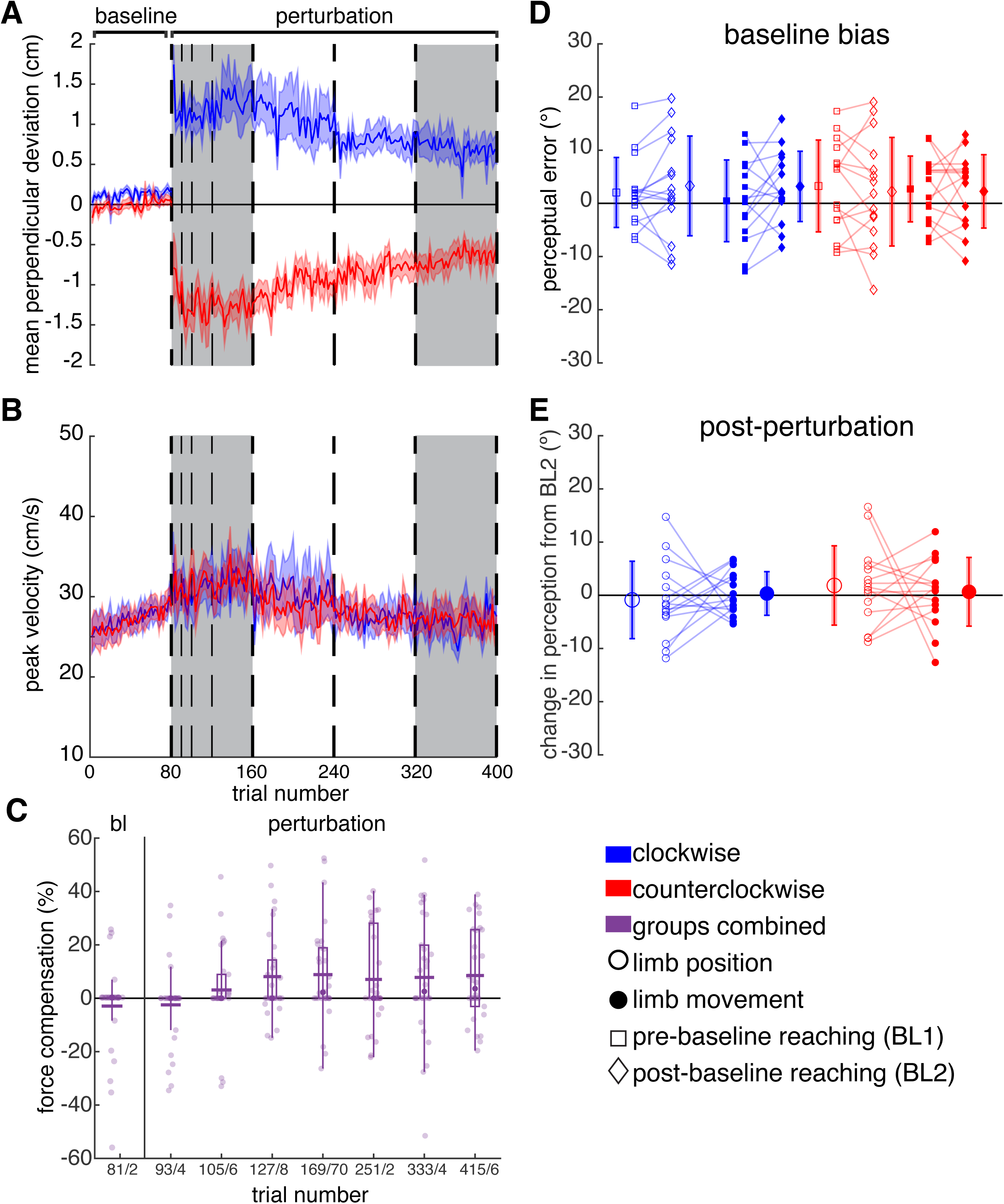
Experiment 2 results. **A-B.** Group mean time series of the mean perpendicular deviation values (**A**) and reach peak velocity (**B**) for each perturbation direction. Note that the same participants performed both perturbation directions with the trials randomly interleaved. Vertical dashed lines depict motor catch trials. Catch trials were performed in pairs, probing each field direction. Thicker dashed lines denote points at which motor catch trials were followed by a 1-2 minute break. Grey shaded regions denote the early and late phases of the perturbation period. **C.** Force compensation on motor catch trials. Vertical boxes indicate the interquartile range, and whiskers indicate the data range excluding outliers. Horizontal lines show the distribution means, while solid markers indicate the median. Transparent markers show subject data. **D.** Perceptual errors exhibited at baseline for both follow-through target trials (mapped to the CW and CCW fields in the perturbation phase of the adaptation task). The report error is shown for each baseline test block within each perceptual probe type for both field directions. **E.** The change in perceptual error between the sensory perception test blocks post-baseline reaching (BL2) and post-perturbation. **D-E.** The smaller markers indicate subject means. Connecting lines depict the within-subject behavior across test phase (**D**) and perceptual probe type (**E**). The larger circle markers indicate group means with error bars indicating the standard deviation.

Figure 3b shows the time series of peak velocity values for reaches in each field direction. Statistical analysis showed a significant effect of phase (*F*(1,2388)=342.890, *p*<.001) that resulted from greater peak velocities in the first, compared to the final, 40 trials of the perturbation phase. Thus, participants generally slowed their reaches over the course of the task. Neither the effect of field (*F*(1,2388)=1.222, *p*=.269) nor the phase-by-field interaction (*F*(1,2388)=.006, *p*=.936) was significant, suggesting no systematic differences in peak velocity between the two perturbation directions. We assessed the correlation between the change in peak velocity and reach MPD over the perturbation phase; it was significant for the CW field (*r*=.784, *p*<.001), but not for the CCW field (*r*=.228, *p*=.413). Overall, this indicates that although participants did tend to slow their reaches over the course of the perturbation, and this may have contributed to reducing reach MPD on CW field trials, this was not the case for all reach trials.

Finally, analysis of force compensation showed a significant effect of time point (*F*(7,218)=2.416, *p*=.021; Fig. 3c). The effect was driven by increasing force compensation values throughout the perturbation phase, mirroring the evidence for adaptation seen in the main analysis. However, post hoc tests showed that none of the pairwise comparisons survived Bonferroni correction.

#### Dual force-field adaptation did not produce evidence of a context-dependent realignment of perception

As in Experiment 1, we first analyzed perceptual reports at baseline to characterize any differences across phases or between follow-through targets (i.e., that would be mapped to a particular force field direction in the perturbation phase; Fig. 3d). The results showed no significant effects, suggesting that perceptual reports were similar before and after baseline reaching and across probe types, and were not differently influenced by cueing with the follow-through target (phase: *F*(1,1418)=.449, *p*=.503, BF_01_=13.812; probe: *F*(1,1418)=.361, p=.548, BF_01_=14.225; follow-through target: *F*(1,1418)=.157, BF_01_=16.196, *p*=.692; phase-by-probe: *F*(1,1418)=.343, *p*=.558, BF_01_=196.679; phase-by-target: *F*(1,1418)=2.307, *p*=.129, BF_01_=216.965; probe-by-target: *F*(1,1418)=.104, *p*=.747, BF_01_=233.706; phase-by-probe-by-target: *F*(1,1418)=.055, *p*=.815, BF_01_=846.141).

To remain consistent with Experiment 1, post-perturbation perceptual reports were normalized by subtracting the mean of the post-baseline perceptual reports (Figure 3e). Statistical analysis showed no significant effects (probe: *F*(1,702.001)=.000, *p*=.997, BF_01_=12.468; field direction: *F*(1,702.001)=1.359, *p*=.244, BF_01_=6.242; probe-by-field: *F*(1,702.001)=.851, *p*=.357, BF_01_=71.490), which suggested that there were no systematic differences in either limb position or movement perception across force field directions following the dual-adaptation task. Additionally, none of the distributions were significantly different from zero, indicating that perception had not systematically shifted from the baseline (limb movement, CCW: *t*(179)=0.529, *p*=.598, BF_01_=10.473, CW: *t*(179)=0.275, *p*=.783, BF_01_=11.578; limb position, CCW: *t*(179)=1.372, *p*=.171, BF_01_=4.779, CW: *t*(179)=-0.632, *p*=.529, BF_01_=9.878).

## DISCUSSION

A growing body of evidence shows that in addition to changing motor commands, adaptation also triggers a realignment of somatosensory perception. The co-occurrence of these two phenomena has led to hypotheses that they are mediated by a common underlying mechanism (Rossi et al. 2021; Tsay et al. 2022), which in turn predicts that motor behavior and somatosensory realignment following adaptation should exhibit similar properties. Here, we tested and found no clear evidence that sensory realignment and motor adaptation following force-field adaptation behave similarly across three properties: whether their magnitudes are correlated (specifically for sense of limb movement versus static position), whether they generalize to reach directions adjacent to the trained one, and whether they display context-dependence.

In the first experiment, participants making reaching movements to a single target at 90 degrees successfully adapted to a single force field that pushed their hand in either a clockwise or counterclockwise direction. We found that following the perturbation phase of the task, somatosensory perception of limb movement to the same target location was significantly biased in the direction of the perturbation. That is, participants who adapted to the clockwise force field tended to perceive passive limb movements to the 90-degree target as being to the right of the actual movement path. In contrast, participants who adapted to the counterclockwise force field tended to perceive the same movement path as biased to the left. Like the adapted movement pattern, the realignment of limb movement perception washed out following a period of reaching without the force field perturbation. Notably, the systematic biasing of perception only occurred when participants were asked to report the perceived movement path following the perturbation phase. When their hands were passively moved to the 90-degree target and held in place, participants showed no systematic bias in somatosensory perception of the position of their index fingertip.

Our study is the first, to our knowledge, to demonstrate differences in the realignment of somatosensory perceptions of limb movement and position following adaptation in the upper extremity. A distinction between these two senses is supported by prior studies of somatosensation (Brown et al. 2003a, 2003b; Proske 2019; Wong et al. 2024), but its nature is poorly understood. Prior work using ischemic block to transiently paralyze and anesthetize the forearm while individuals attempted wrist flexion and extension movements showed that somatosensory perception of the movements persisted even without muscle activity or peripheral feedback (Walsh et al. 2010). A similar study that used a local paralytic agent to block muscle activity while preserving peripheral feedback also showed that the perception of wrist movement persisted despite conflicting afferent information (Smith et al. 2009). Thus, the conscious perception of limb movement does not fully depend on peripheral feedback signals. Instead, it may use information derived from motor commands, such as sensory or cognitive predictions (Bhanpuri et al. 2013; Weeks et al. 2017); although, as will be discussed, these predictions may not guide perception in quite the same way as adaptation. In contrast, somatosensory perception of static limb position may rely more heavily on peripheral feedback. Indeed, we have previously found that perceptions of static position are more susceptible to a visual-somatosensory conflict than perceptions of movement (Wong et al. 2024).

Interestingly, the selective biasing of limb movement perception we observed did align with the nature of the force field perturbation, which affected the execution of reaching movements but did not misalign visual and somatosensory feedback of the index fingertip position during them. Our findings dovetail with studies of somatosensory realignment following adaptation to analogous dynamic perturbations in the lower extremity (Sombric et al. 2019; Vazquez et al. 2015) and suggest that the somatosensory realignment previously observed following force field adaptation may have reflected changes in the perception of movement rather than position (Haith et al. 2008; Mattar et al. 2013; Ohashi et al. 2019; Ostry et al. 2010). It would be interesting for future work to test whether the nature of somatosensory realignment following adaptation in other paradigms also displays coherence with the nature of the perturbation used. For example, visuomotor perturbations involve both a recalibration of sensorimotor predictions and multisensory integration. It is possible that with the coactivation of adaptation and multisensory integration, somatosensory realignment in visuomotor paradigms might result from a biasing of both the perceptions of position and movement (Rossi et al. 2021).

Despite this coherence between the observed realignment and the nature of the force field perturbation, however, and in spite of our effort to carefully dissociate changes in sense of limb movement and position, we observed no correlation between the extent of sensory realignment and motor adaptation. This result differed from prior studies of somatosensory realignment following force field adaptation (Mattar et al. 2013; Ohashi et al. 2019). It is possible that this discrepancy results from differences between our method of probing somatosensory perception (e.g., dissociating sense of limb movement and position) and that used in prior work. Specifically, prior studies that found a correlation between force field adaptation and somatosensory realignment tested general somatosensory perception using a two-alternative forced choice task that had participants judge whether a passive displacement had moved their dominant hand to the right or left of a visual stimulus (Mattar et al. 2013; Ohashi et al. 2019). Using a standard psychophysical design, several displacements were tested, and perception was quantified as the bias of a psychometric curve fit to the report data. The result was a broad measure of somatosensory perception that may have combined the sense of limb movement and position, but in contrast, was likely more robust to trial-by-trial variability than the method used in our study. Indeed, our study design involved a relatively small number of trial repetitions for each sensory perception probe. It is possible that with an increased number of repetitions per subject, we would have seen evidence of correlation. Yet, it is worth noting that across adaptation paradigms, the overall picture is mixed. Studies of somatosensory realignment following visuomotor adaptation often fail to find correlations between sensory and motor measurements (Barkley et al. 2014; Cressman et al. 2010; Cressman and Henriques 2009; Modchalingam et al. 2019; Nourouzpour et al. 2015; Salomonczyk et al. 2011). A lack of correlation was also noted for somatosensory realignment in the lower extremity following adaptation to a dynamic perturbation during gait (Rossi et al. 2019). Inconsistent findings of correlations between somatosensory realignment and adaptation across studies argue against the two phenomena arising from a shared mechanism and the need for more precise quantification of changes in perceived limb movement versus position.

A second piece of evidence against a unified mechanism underlying both adaptation and somatosensory realignment is the observed lack of generalization for sensory realignment. Force field adaptation is known to show a cosine-like generalization pattern, in which the adapted movement pattern gradually decays as reach directions move further away from the one trained. Prior work has shown the adaptation generalization window to be within about 45 degrees of the trained movement; however, the effect can be asymmetric across clockwise and counterclockwise directions (Donchin et al. 2003; Gandolfo et al. 1996; Gonzalez Castro et al. 2011; Huang and Shadmehr 2007; Mattar and Ostry 2007; Rezazadeh and Berniker 2019; Sainburg et al. 1999; Thoroughman and Shadmehr 2000). We found that the realignment of limb movement perception did not systematically generalize to movement directions rotated just 10 degrees clockwise and counterclockwise from the one performed in the adaptation task. Although we did not directly assess generalization to the 80 and 100 degree directions in the motor domain, the generalization targets were chosen to be within the range observed in previous studies of force field adaptation where generalization is expected to be as great as 96% (Gonzalez Castro et al. 2011). Analysis of perceptual biases at baseline suggest that the 10-degree separation of targets fell within participants’ report variance, which may have limited our capacity to interpret the generalization results. Nonetheless, selective perceptual realignment at the actual trained movement path suggests some distinction between it and the generalization directions. Our findings are also consistent with prior work studying somatosensory realignment following adaptation to a visuomotor perturbation, which found that the pattern of generalization differed from that observed in the motor domain (Cressman and Henriques 2015). While further study is needed, these results suggest that somatosensory realignment may not generalize in the same manner as motor adaptation.

Our second experiment examined another property of adaptation: context-dependence. It is well documented that individuals can simultaneously adapt to two opposing perturbations, so long as each is paired with a distinct sensory and/or motor context cue (Addou et al. 2011; Cothros et al. 2009; Howard et al. 2013, 2015; Ingram et al. 2013; Osu et al. 2004; Richter et al. 2004; Sheahan et al. 2016; Wada et al. 2003). Following dual adaptation, the presentation of each context cue elicits opposing after-effects, indicating that the formation and recall of sensorimotor calibrations are context dependent. We tested whether post-adaptation somatosensory realignment also shows context-dependence by comparing participants’ perceptions of limb movement and position following presentation of the context cue before and after a dual adaptation task. Our results showed no evidence of context-dependent somatosensory realignment.

No evidence of realigned limb position perception following dual adaptation aligns with the results of Experiment 1. However, the lack of systematic bias in limb movement perception in Experiment 2 contrasts with the outcome of our single-field experiment. Given that somatosensory realignment lags behind motor adaptation (Mattar et al. 2013), and motor adaptation tends to be slower in dual-compared to single-perturbation paradigms, it is possible that evidence for context-dependent realignment would appear with longer task durations. Additionally, the difference in results may have stemmed from differences in reach kinematics owing to different perturbation distances across the two adaptation tasks: 20 cm in Experiment 1 versus 12 cm in Experiment 2. However, another interpretation is that the evidence here favoring the null hypothesis represent a further distinction between adaptation and somatosensory realignment. While it is possible for the sensorimotor system to maintain and rapidly call upon opposing calibrations for motor output, the effects may cancel out at the level of sensory perception. The capacity to rapidly adjust motor output is advantageous, as it helps maintain movement accuracy in the face of environmental changes, but equally rapid changes to sensory perception would likely be cognitively destabilizing.

When taken together, the results of both experiments align with prior literature suggesting that sensory perception can be mediated independently of sensory function for motor control. Sensory perception is well known to be susceptible to illusions that fail to bias motor output. For example, in the visual domain, the Muller-Lyer illusion is a robust perceptual phenomenon in which individuals perceive line segments as different lengths depending on the direction of arrowheads placed at either end (Mack et al. 1985). Despite the illusion, reaching movements directed to the endpoints of each segment are generally accurate (Bernardis et al. 2005; Binsted and Elliott 1999; Gentilucci et al. 1997; Thompson and Westwood 2007). Visual-tactile illusions, such as the Rubber Hand illusion, in which simultaneous visual and tactile input induces the perceived embodiment of a fake hand (Botvinick and Cohen 1998), also fail to induce motor errors on subsequent reaching movements (Kammers et al. 2006, 2009). While the precise root of this distinction remains unclear, it is likely that sensory perception is subject to cognitive processing, especially when it is tested using individuals’ conscious reports, as is often the case. Conversely, the utilization of sensory information for motor control may be less permeable to this additional cognitive input.

In conclusion, this study examined properties of somatosensory realignment following force field adaptation in two experiments. The first experiment replicated prior results showing that adaptation in this paradigm coincides with a realignment of somatosensory perception in the direction of the force perturbation. We additionally found that the realignment reflected a selective biasing of the perception of movement, and not of static position. However, the magnitude of realignment did not correlate with the magnitude of adaptation, and the realignment of limb movement perception did not generalize to reach directions flanking the one performed in the adaptation task. In a second experiment, we found that dual adaptation to opposing force fields did not result in context-dependent realignment of limb movement perception. Overall, our results suggest that although the nature of somatosensory realignment following adaptation might be coupled to that of the perturbation, the phenomena may not arise from the same underlying mechanism.

### Experimental data and custom analysis codes are available here

https://osf.io/rj2ug/overview

## Acknowledgements

We thank members of the Movement Science Focus Area at the Jefferson Moss Rehabilitation Research Institute for helpful discussions. This work was supported by pilot funding from the Jefferson Moss Rehabilitation Research Institute Peer Review Committee and NSF grant BCS2444305.

## Author Contributions

AST and ALW designed the experiments. AST programmed the experimental software. AST and DZ collected experimental data and performed the data analysis. AST and ALW interpreted the experimental results and wrote the manuscript.

## Notes

https://osf.io/rj2ug/overview

### Competing Interest Statement

The authors have declared no competing interest.

### Summary of Updates

1. Materials originally contained in a Supplement are now included in the main text. 2. Bayes Factors are included for null results from the analysis of perceptual reports. 3. Figures 2 and 3 have been updated to show baseline perceptual biases and within-subject effects. 4. The introduction and discussion have been updated to more clearly justify the methods, explain the significance of the findings, and the limitations of the study.

